# Enhanced Wound Healing and Anti-Inflammatory Effects of Vitamin C-Loaded Hyaluronic Acid–Collagen Scaffolds in Preclinical Rat Models

**DOI:** 10.1101/2025.08.06.665568

**Authors:** Ahmed Salem, Samah El-Ghlban, A.S. Montaser, Mohamed F. Abdelhameed, Mohamed F. Attia

## Abstract

Wound healing is a complex biological process critical for restoring skin integrity after injury. However, chronic wounds present significant clinical challenges due to persistent inflammation, disrupted collagen synthesis, and susceptibility to infection. Bioactive scaffolds have emerged as promising therapeutic strategies to enhance tissue regeneration by modulating cellular behavior and extracellular matrix (ECM) dynamics. This study explores a hyaluronic acid-collagen (HyCol) scaffold enriched with vitamin C (VC), producing (VC-HyCol) to improve wound healing in preclinical rat models. Hyaluronic acid and collagen, key ECM components, provide structural and biochemical support, while vitamin C acts as both a collagen biosynthesis cofactor and an antioxidant to counteract oxidative stress. The scaffold was designed to emulate the native ECM microenvironment, facilitating fibroblast proliferation, keratinocyte migration, and angiogenesis. Physicochemical characterization, biocompatibility assessments, and *in vivo* wound healing experiments were performed to evaluate its therapeutic efficacy. Results demonstrated that the incorporation of vitamin C significantly enhanced fibroblast activity, reduced inflammatory markers, and accelerated tissue regeneration compared to control groups. Histological and molecular analyses further confirmed enhanced collagen deposition and neovascularization, indicating faster and more organized wound repair. These findings highlight the potential of this multifunctional scaffold as an advanced wound dressing, with significant implications for regenerative medicine and clinical wound management.

**Graphical Abstract:** 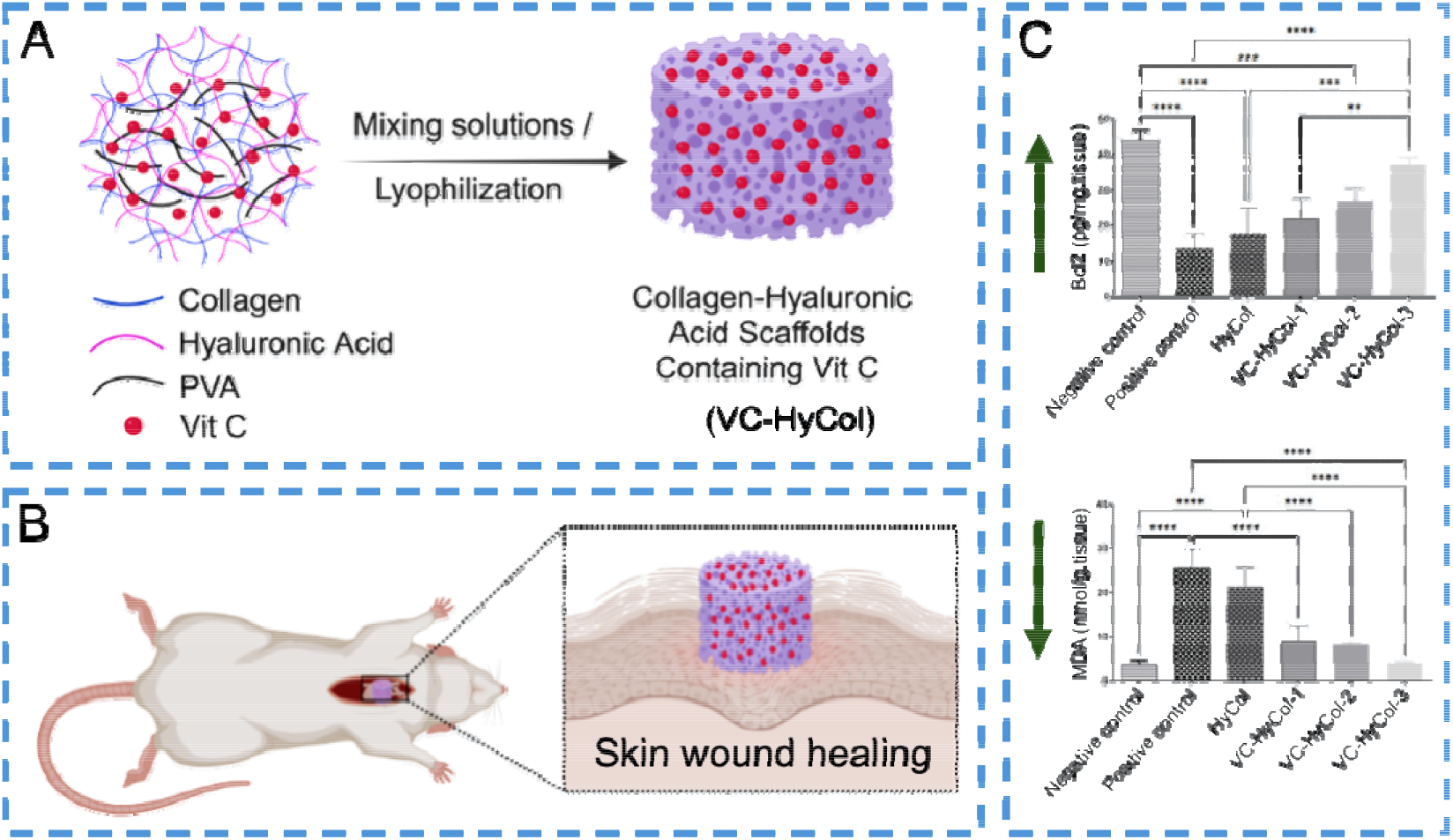

## Introduction

Chronic wounds represent a significant clinical burden, affecting millions of patients worldwide and posing substantial challenges to healthcare systems [1-4]. Unlike acute wounds, which follow a well-orchestrated healing process involving hemostasis, inflammation, proliferation, and remodeling, chronic wounds are characterized by prolonged inflammation, impaired extracellular matrix (ECM) deposition, and delayed re-epithelialization [5-8]. Factors such as diabetes, vascular insufficiency, and persistent infections exacerbate these complications, leading to non-healing ulcers that are resistant to conventional therapies [9, 10]. Traditional wound dressings, while providing passive protection, often fail to address the underlying biological dysregulation [11, 12]. Consequently, there is an urgent need for advanced therapeutic strategies that actively promote tissue regeneration by modulating cellular behavior and restoring the wound microenvironment [13, 14].

Tissue engineering has emerged as a promising approach to overcome these limitations by developing bioactive scaffolds that mimic the native ECM and deliver therapeutic agents to enhance healing [15] xx. An ideal scaffold should provide structural support, facilitate cell adhesion and migration, and regulate key biological processes such as angiogenesis, collagen synthesis, and inflammation [16]. Among the various biomaterials explored, hyaluronic acid (HA) and collagen have gained considerable attention due to their biocompatibility, biodegradability, and critical roles in wound repair [17]. HA, a glycosaminoglycan, enhances tissue hydration and cell motility [18], while collagen, the primary structural protein in the skin, provides tensile strength and serves as a scaffold for fibroblast infiltration [19]. Combining these biomaterials can synergistically improve mechanical stability and bioactivity, making them suitable for wound healing applications [20, 21].

Despite their many advantages, HA-collagen scaffolds alone may not sufficiently address the oxidative stress and inflammatory dysregulation that impede chronic wound healing [22]. In such wounds, excessive accumulation of reactive oxygen species (ROS) leads to cellular damage and sustained inflammation, ultimately delaying tissue repair [23]. To counteract this, antioxidant supplementation has emerged as a promising strategy to reduce oxidative injury and promote regeneration [24]. Vitamin C (ascorbic acid), a well-known antioxidant, plays a dual role in this context: it neutralizes ROS and serves as a critical cofactor in collagen biosynthesis by stabilizing the hydroxylation of proline and lysine residues in procollagen [25]. However, its clinical application is limited by its instability in aqueous environments, which reduces its bioavailability and therapeutic potential [26]. Incorporating vitamin C into scaffold-based delivery systems offers a viable solution, enabling controlled release and prolonged activity at the wound site. This approach not only enhances local antioxidant defense but also supports sustained collagen production and tissue remodeling [26].

This study introduces a novel macroporous hyaluronic acid-collagen (HyCol) scaffold loaded with vitamin C (VC-HyCol) to accelerate wound healing through combined structural, biochemical, and antioxidant mechanisms. The scaffold was engineered to exhibit an interconnected porous architecture, facilitating nutrient diffusion, cell infiltration, and vascular ingrowth. Vitamin C was incorporated at varying concentrations to optimize its release kinetics and therapeutic effects. We hypothesized that this multifunctional scaffold would enhance fibroblast proliferation, reduce oxidative stress, and modulate inflammatory mediators, thereby promoting faster and more organized tissue regeneration.

To evaluate this hypothesis, we conducted comprehensive physicochemical characterization, biocompatibility assessments, and in vivo wound healing experiments in a preclinical rat model. The scaffold’s microstructure, chemical composition, mechanical properties, and degradation profile were analyzed to ensure optimal performance in a dynamic wound environment. *In vitro* studies assessed fibroblast viability and antimicrobial activity, while *in vivo* experiments examined wound closure rates, histopathological changes, and molecular markers of inflammation and angiogenesis. Our findings demonstrate that the VC-HyCol-3 scaffold with higher concentration of VC significantly improves healing outcomes compared to conventional treatments, highlighting its potential as an advanced wound dressing for regenerative medicine. By integrating HA and collagen with vitamin C, this study advances the field of bioactive wound dressings, offering a clinically translatable solution for chronic wound management. The scaffold’s ability to simultaneously provide structural support, enhance ECM deposition, and counteract oxidative stress positions it as a promising therapeutic platform. Future research could explore its applicability in diabetic ulcers, burn injuries, and other hard-to-heal wounds, further validating its role in next-generation wound care.

## Experimental section

### Chemicals and Reagents

The study utilized Type I collagen (bovine origin) and polyvinyl alcohol (PVA, MW 31–50 kDa) from Sigma-Aldrich, along with high-molecular-weight hyaluronic acid (HA, 1.5-1.8 kDa) from Lifecore Biomedical. Ethanol (70-100%) was obtained from Fisher Scientific, while Vitamin C (VC) and other cell culture reagents, including Dulbecco’s Modified Eagle Medium (DMEM), penicillin-streptomycin, trypsin-EDTA (0.25%), fetal bovine serum (FBS), and trypan blue, were sourced from Gibco and Sigma-Aldrich. For cytotoxicity assessment, human skin fibroblasts (HSF) were used, along with 3-(4,5-dimethylthiazol-2-yl)-2,5-diphenyltetrazolium bromide (MTT) and dimethyl sulfoxide (DMSO) from Sigma-Aldrich. Histological processing involved neutral buffered formalin (Sigma-Aldrich) and hematoxylin/eosin (H&E) staining reagents (Thermo Scientific/Sigma-Aldrich). Immunohistochemistry was performed using citrate buffer (BioGenex), a Vectastain ABC-HRP kit (Vector Laboratories), 3,3’-diaminobenzidine (DAB, Sigma-Aldrich), and an anti-TGF-β primary antibody (Servicebio, GB11179). Protein analysis was conducted using RIPA lysis buffer (Sigma-Aldrich), PVDF membranes (Millipore), and enhanced chemiluminescence substrate (Thermo Scientific). Oxidative stress markers were evaluated using MDA (Biovision K739-100) and GSH (Biovision K464-100) assay kits, along with thiobarbituric acid (TBA, Sigma-Aldrich, Cat# T5500) and 5,5’-dithiobis-(2-nitrobenzoic acid) (DTNB, Sigma-Aldrich, Cat# D8130). Cytokine levels were measured via ELISA kits for IL-1β (SL0402Ra), TNF-α (SL0722Ra), VEGF (SL0740Ra), CD34 (SL1701Ra), and Bcl-2 (SL0108Ra) from Sunlong Biotech, with TMB substrate (Sigma-Aldrich, Cat# T0440) and 2N H□SO□ (Sigma-Aldrich) for signal development. For antimicrobial testing, nutrient agar (BD Difco, Cat# 213000), potato dextrose agar (Sigma-Aldrich, Cat# P2182), and sterile saline (Thermo Fisher, Cat# J66984) were used. Reference strains (*S. aureus* ATCC 6538-P, *E. coli* ATCC 25933, *C. albicans* ATCC 10231, *A. niger* NRRL-A326) were obtained from ATCC and USDA-ARS, with sterile textile discs (Whatman, Cat# 2017-915) serving as scaffold carriers. All media were sterilized by autoclaving (121 °C, 15 min), and microbial densities were confirmed via colony counting.

## Methods

### Preparation of macroporous hyaluronic acid-collagen (HyCol) hybrid scaffolds

Macroporous hyaluronic acid-collagen (HyCol) hybrid scaffolds were fabricated by blending sterile-filtered Type I collagen (3 wt%, bovine origin) with high-molecular-weight hyaluronic acid (HA, 3 wt%, 1.5–1.8 kDa) in a 1:1 volume ratio. To enhance structural integrity, polyvinyl alcohol (PVA, 5 wt%) was prepared in deionized water at 80 °C for 3 h under a nitrogen atmosphere, followed by sequential filtration (5 μm → 0.22 μm) to ensure sterility and homogeneity. Vitamin C (VC) was incorporated at three different concentrations: 0.5 g (0.74% w/v), 1 g (1.48% w/v), and 1.5 g (2.22% w/v), dissolved in 10 mL of PBS (pH 7.4) immediately before use. The VC solution was added dropwise (1 mL/min) to the collagen, followed by the incorporation of HA. Crosslinking was then initiated by adding 7.5 mL of the prepared PVA solution. The resulting hydrogel was cast into 12-well plates and underwent controlled thermal phase separation (4 °C/2 h, −20 °C/12 h, −80 °C/4 h). The samples were lyophilized (0.05 mbar, 72 h). The resulting macroporous scaffolds were stored in vacuum-sealed bags until further use. All scaffold compositions and ratios are represented in **Table 1**.

**Table 1.** summarizes the composition of the prepared scaffolds, including the weights of all components and the vitamin C (VC) loading capacity (LC).

| # | Scaffold | VC (g) | Col (g) | HA (g) | PVA (mg) | LC% |
| --- | --- | --- | --- | --- | --- | --- |
| 1 | HyCol | 0 | 0.9 | 0.9 | 37.5 | 0 |
| 2 | VC-HyCol-1 | 0.5 | 0.9 | 0.9 | 37.5 | 21.39 |
| 3 | VC-HyCol-2 | 1 | 0.9 | 0.9 | 37.5 | 35.24 |
| 4 | VC-HyCol-3 | 1.5 | 0.9 | 0.9 | 37.5 | 44.94 |

The LC was calculated using the following equation:

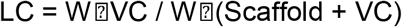

Where, W□VC = weight of loaded vitamin C and W□(Scaffold + VC) = total weight of the scaffold and Vitamin C.

### Scaffold Characterization

#### Attenuated Total Reflectance Fourier-Transform Infrared Spectroscopy (ATR-FTIR)

Chemical composition and crosslinking efficacy were assessed using ATR-FTIR (Nicolet iS50, Thermo Scientific). Triplicate measurements of dried scaffolds were performed (32 scans, 4 cm□^1^ resolution) across the mid-infrared range (4000–400 cm□^1^). Acquired spectra were baseline-corrected (OMNIC v9.0) and deconvoluted via Gaussian-Lorentzian curve fitting (R^2^ > 0.995) to evaluate collagen crosslinking, hyaluronic acid esterification, and vitamin C stability.

#### Scanning Electron Microscopy (SEM) analysis

Scaffold morphology was examined using an environmental SEM (Quanta FEG 250, FEI Company) operated at 10 kV accelerating voltage under high vacuum (≤1 × 10□^3^ Pa). Samples were mounted on aluminum stubs with conductive carbon tape and imaged without metallic coating, utilizing the instrument’s native charge compensation capabilities.

#### Swelling behavior analysis

Lyophilized scaffolds (n=5 per group) with precisely measured dry weights (Wd, ±0.01 mg) were immersed in PBS (pH 7.4) at 37 °C. At specified time intervals (0.5, 1, 2, 4, 8, 12, and 24 h), samples were removed, surface moisture was blotted using lint-free filter paper, and wet weights (Ws) were recorded. The swelling ratio was calculated as:

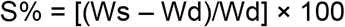

#### Porosity quantification

Scaffold porosity was determined using an ethanol displacement method with absolute ethanol (99.8% purity, ρ = 0.789 g/cm^3^ at 25 °C). Initial ethanol volume (V□ = 10.00 ± 0.05 mL) was measured in a Class A graduated cylinder (±0.1 mL accuracy). Dry scaffolds (Wd) were immersed in ethanol and subjected to vacuum degassing (25 inHg, 3 cycles of 1 min ON/OFF), with the saturated volume (V□) recorded after 15 min equilibration. After scaffold removal and 30 s drainage at 45°, residual ethanol volume (V□) was measured. Porosity was calculated as:

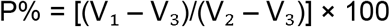

### Antimicrobial Activity Evaluation

The antimicrobial properties of the engineered scaffolds were assessed against representative pathogenic strains using the disc diffusion assay [27]. The test microorganisms included both bacterial and fungal species: Gram-positive Staphylococcus aureus (ATCC 6538-P), Gram-negative Escherichia coli (ATCC 25933), the yeast Candida albicans (ATCC 10231), and the filamentous fungus Aspergillus niger (NRRL-A326). For bacterial and yeast testing, nutrient agar plates were uniformly inoculated with 0.1 mL of microbial suspension (10^5^–10^6^ CFU/mL), while potato dextrose agar plates were employed for fungal assessments using a standardized spore suspension (0.1 mL of 10^6^ spores/mL). Sterile scaffold-loaded textile discs (15 mm diameter) were aseptically placed on the inoculated surfaces. To enhance compound diffusion, plates were pre-incubated at 4 °C for 2–4 h before transfer to optimal growth conditions: 37 °C for 24 h for bacterial strains and 30 °C for 48 h for fungal/yeast cultures, maintained in an upright position to prevent condensation interference. Following incubation, antimicrobial efficacy was quantitatively determined by measuring the diameter of inhibition zones (in millimeters) surrounding each test disc.

### Cell Viability Assessment (MTT Assay)

Cell viability was assessed using the MTT assay [28]. Human dermal fibroblasts were seeded into 96-well plates at a density of 5,000 cells per well in 100□μL of complete DMEM and incubated for 24 h at 37□°C in a 5% CO_2_ atmosphere to allow cell attachment. After this period, test scaffolds, prepared in serum-free medium, were added at concentrations ranging from 1, 2.5, and 5□mg per well (100□μL) and incubated for an additional 24 h. The medium was then replaced with 20□μL of MTT solution (1□mg/mL in PBS), and cells were incubated for 4 h at 37□°C. Formazan crystals formed during this process were solubilized in 100□μL of absolute DMSO under gentle shaking for 30 min at room temperature (RT). Absorbance was recorded at 570□nm with a reference wavelength of 690□nm using a FLUOstar Omega microplate reader (BMG Labtech, Germany). Cell viability was calculated relative to untreated controls, with all conditions tested in triplicate and inter-assay variability kept below 15%.

### Animal Study Protocol

Thirty adult male Sprague-Dawley rats (150–200 g) were procured from the National Research Centre’s animal facility. The animals were kept for one week under standard conditions (24–27 °C, 12-h light/dark cycle). At the Menofia University Faculty of Science, rats were kept in custom plastic rodent cages. In each cage, there were five rats. Rats were given unlimited access to food and water. Rats were randomized into six groups (n=5/group): Group I (sham control (negative control), intact skin), Group II (positive control, untreated full-thickness wounds), Group III (wounds treated with HyCol scaffold), and Groups IV, V, and VI (wounds treated with VC-HyCol-1, VC-HyCol-2, and VC-HyCol-3, respectively). Under ketamine/xylazine anesthesia (75/8 mg/kg, i.p.), dorsal fur was clipped and disinfected, and circular full-thickness excisional wounds (100–125 mm^2^) were created aseptically using surgical blades and forceps. Scaffolds were applied daily under brief isoflurane sedation (2%). Wound progression was monitored via standardized photography at days 0, 7, 10, and 14 post-wounding, with planimetric analysis (ImageJ) used to calculate percentage closure: [(A□ − A□)/A□] × 100, where A□ = initial area and A□ = area at timepoint. Terminal procedures included tissue collection for histopathological and molecular analyses. The study complied with ARRIVE guidelines [29] and was approved by Menoufia University’s IACUC (MUFS/F/Bio/1/25).

### Oxidative Stress and Inflammatory Marker Analysis

Oxidative stress markers were evaluated using standard biochemical techniques [30]. Malondialdehyde (MDA) levels were quantified via the thiobarbituric acid reactive substances (TBARS) assay, in which tissue homogenates were incubated with 0.67% thiobarbituric acid at 95□°C for 60 min, and absorbance was subsequently measured at 532□nm. Reduced glutathione (GSH) was assessed using an enzymatic recycling method involving 5,5′-dithiobis(2-nitrobenzoic acid) (DTNB), with kinetic readings taken at 412□nm. Inflammatory and angiogenic mediators, including interleukin-1β (IL-1β), tumor necrosis factor-α (TNF-α), vascular endothelial growth factor (VEGF), CD34, and Bcl-2, were quantified using sandwich ELISA kits following the manufacturers’ protocols [31-33]. Briefly, pre-coated 96-well plates were incubated with 100□μL of sample per well for 2 h at RT, followed by detection antibody incubation for 1 h and development with tetramethylbenzidine (TMB) substrate for 15 min. The reaction was terminated with 2N sulfuric acid, and absorbance was recorded at 450□nm with a reference at 570□nm. All measurements were performed in triplicate, with standard curves (R^2^ > 0.98) and appropriate controls to ensure assay reliability.

### Western Blot Analysis

Protein expression was analyzed using an optimized Western blot protocol [34]. Tissue samples were lysed in ice-cold RIPA buffer (50□mM Tris-HCl, pH 8.0; 150□mM NaCl; 1% NP-40; 0.5% sodium deoxycholate; 0.1% SDS) supplemented with a protease and phosphatase inhibitor cocktail (cOmplete™, Roche). Homogenates were incubated for 30 min at 4□°C with continuous agitation, followed by sonication (three 15-second pulses at 30% amplitude) and centrifugation at 16,000×g for 20 min at 4□°C to remove debris. Protein concentrations were determined using the BCA assay (Pierc™), with bovine serum albumin as the standard. Equal amounts of protein (30□μg per lane) were denatured and resolved on 15% SDS-PAGE gels using a two-step voltage protocol (90□V through the stacking gel, then 120□V through the resolving gel). Proteins were transferred to 0.2□μm PVDF membranes via semi-dry transfer (100□V, 90 min, 4□°C) using standard Tris-glycine-methanol buffer. Membranes were blocked in 5% non-fat dry milk in TBST (20□mM Tris, 150□mM NaCl, 0.1% Tween-20, pH 7.5) for 1 h at RT, then incubated overnight at 4□°C with primary antibodies against NF-κB p65 (1:1000), GAPDH (1:1000), and β-actin (1:3000) (Invitrogen, Mouse species). After washing, membranes were incubated with HRP-conjugated secondary antibodies (1:3000) for 1 h at RT. Protein bands were visualized using enhanced chemiluminescence (SuperSigna™ West Pico PLUS, Thermo Scientific) and imaged with a ChemiDo™ MP system (Bio-Rad). Densitometric analysis was performed using Image La™ software (v6.1, Bio-Rad), with normalization to β-actin or GAPDH and background correction. Each experiment included technical triplicates, positive controls (rat spleen lysate), negative controls (no primary antibody), and molecular weight markers (Precision Plus Protei™ Kaleidoscop™, Bio-Rad) to ensure data integrity. Quantitative assessments of the relative intensity of the blots were analyzed using ImageJ.

### Histopathological Analysis

For optimal morphological preservation, tissue specimens were fixed in 10% neutral buffered formalin (pH 7.4) at RT for 48 h. After fixation, the samples were trimmed to standardized dimensions (5 × 5 mm) and rinsed thoroughly in PBS (pH 7.2) for 24 h. Dehydration was achieved through a graded ethanol series (70%, 80%, 95%, and 100%), with each concentration applied for 1 h. The tissues were then cleared in three changes of xylene (30 min each) and infiltrated with molten paraffin wax (56–58°C) under vacuum for 2 h before embedding. Serial sections of 4–6 μm thickness were cut using a rotary microtome fitted with high-profile blades. To prevent folding, sections were floated on a 40 °C water bath containing 0.1% gelatin and mounted on poly-L-lysine-coated slides. After drying overnight at 37°C, automated staining was performed using Harris hematoxylin (5 min) and eosin Y (1 min). Differentiation was carried out in 0.5% acid alcohol (30 sec), followed by bluing in 0.2% ammonium hydroxide (1 min). Finally, the sections were dehydrated through graded alcohols, cleared in xylene, and mounted with DPX. Slides were examined under a light microscope with digital imaging capabilities, enabling quantitative assessment of epidermal thickness and granulation tissue formation at standardized magnifications (20× for tissue architecture, 40× oil immersion for cellular details). Method validation included control slides in each staining batch and microtome calibration to ensure section thickness consistency (≤5% variation). All procedures adhered to established histological standards (Bancroft & Gamble, 2008) and institutional SOPs (NRC Histopathology Lab, 2023).

### Immunohistochemical Analysis

Immunohistochemical analysis was conducted on paraffin-embedded tissue sections (4–6□μm) mounted on positively charged glass slides. Following deparaffinization and rehydration, antigen retrieval was performed by heating the sections in citrate buffer (pH 6.0) at 95□°C for 20 min. Endogenous peroxidase activity was quenched using 3% hydrogen peroxide for 15 minutes at room temperature. To minimize non-specific binding, slides were incubated with 5% normal goat serum (Vector Laboratories) for 1 hour. Sections were then incubated overnight at 4□°C in a humidified chamber with a rabbit polyclonal anti-TGF-β primary antibody (Servicebio, GB11179; 1:800 dilution). Detection was carried out using the Vectastain ABC-HRP system (Vector Laboratories), involving sequential 30-min incubations with a biotinylated secondary antibody and the avidin-biotin complex. Visualization was achieved using 3,3′-diaminobenzidine (DAB; Sigma-Aldrich) as the chromogen, with staining monitored microscopically to ensure optimal signal development. Negative controls were included by substituting the primary antibody with non-immune serum to confirm specificity. Slides were counterstained with Mayer’s hematoxylin, dehydrated through graded ethanol, cleared in xylene, and mounted using DPX medium. Digital images were captured from ten representative fields per sample at 400× magnification using an Olympus BX-63 microscope equipped with a high-resolution camera. Quantitative analysis of DAB-positive staining was performed using ImageJ software (version 1.53t, NIH), applying threshold-based segmentation to calculate the percentage of stained area. All evaluations were conducted in a blinded manner. Method consistency was ensured through the use of internal positive controls and inter-assay calibration with reference standards.

## Results and discussion

### Scaffold Fabrication and Structural Characterization

Chronic wound healing remains a significant clinical challenge, with conventional scaffolds often failing to address key pathological factors, including prolonged inflammation, oxidative stress, and insufficient extracellular matrix (ECM) regeneration. Existing collagen-based dressings frequently exhibit poor mechanical stability or rapid degradation, while synthetic alternatives lack the bioactive cues necessary for proper tissue remodeling. To overcome these limitations, we developed a vitamin C-loaded hyaluronic acid-collagen (VC-HyCol) hybrid scaffold designed to simultaneously provide structural support, modulate oxidative damage, and accelerate healing through controlled bioactive delivery.

The scaffold’s rigorous fabrication process (**Figure 1A**) was engineered to optimize both biological function and structural integrity. By blending collagen (Col) with hyaluronic acid (Hy) in a 1:1 ratio, we created a biomimetic ECM base that retains natural cell-adhesion motifs while leveraging Hy’s hygroscopic and immunomodulatory properties. Polyvinyl alcohol was incorporated as a thermal-stable crosslinker, with nitrogen-purged synthesis preventing oxidative degradation of sensitive components during the 80 °C dissolution phase. Vitamin C was introduced under argon protection to maintain its redox-active state, ensuring full retention of its antioxidant and pro-collagen synthesis activities. The schematic highlights the sequential mixing, crosslinking, and lyophilization steps, which are critical for preserving the structural integrity and porosity of the final scaffold.

**Figure 1.**
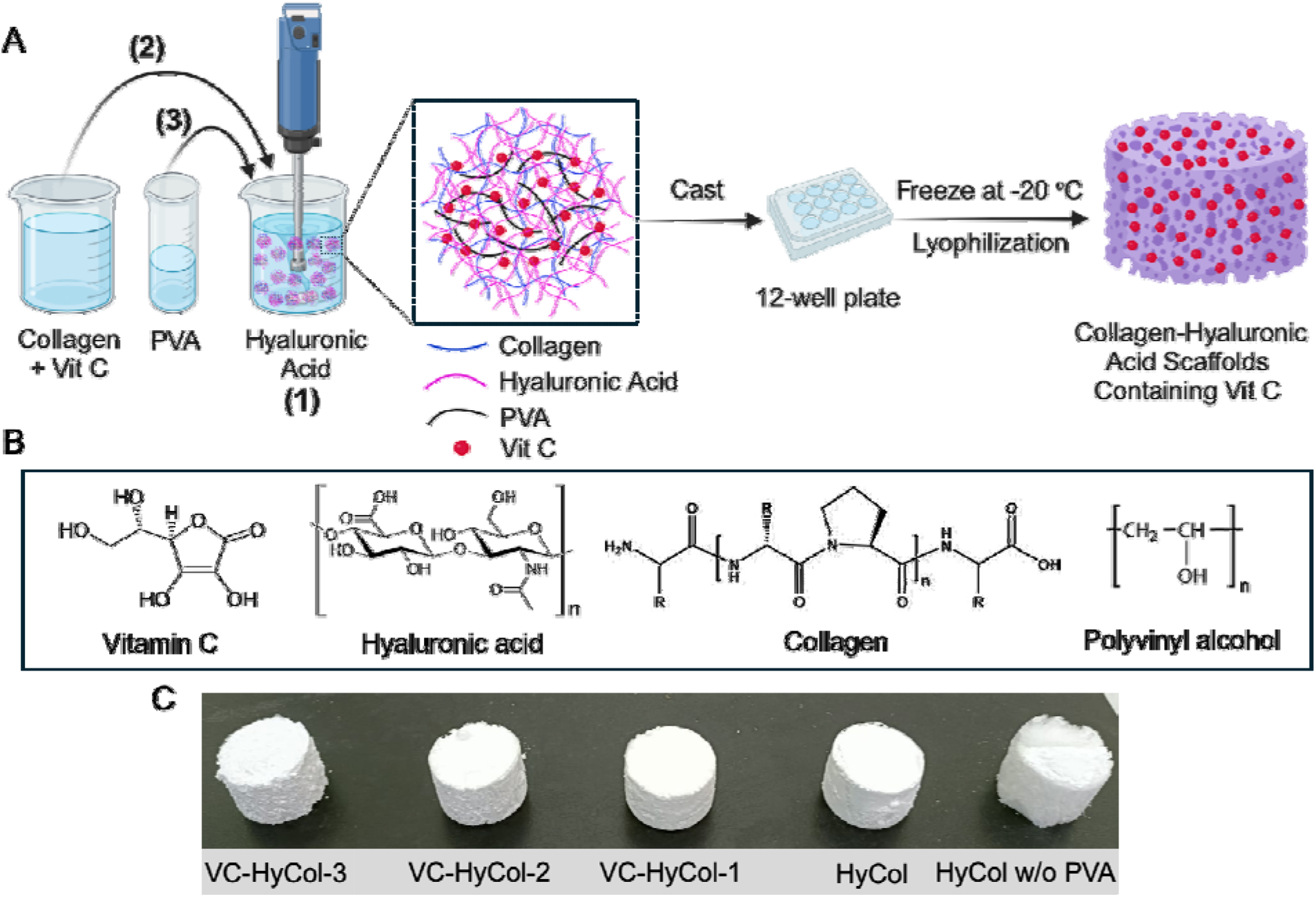
(A) Schematic representation of one-pot synthesis of a series of scaffolds. (B) Chemical structures of all compounds involved in the formulated scaffolds. (C) Digital images of the prepared scaffolds after lyophilization process.

**Figure 1B** illustrates the chemical structures of the key components of the scaffold. Hyaluronic acid and collagen mimic the native extracellular matrix, while vitamin C supports collagen synthesis and provides antioxidant protection, highlighting their suitability for wound healing applications. As shown in **Figure 1C**, the lyophilized VC-HyCol scaffolds exhibit a uniform, sponge-like morphology with well-preserved macroporous architecture across all vitamin C concentrations. This consistent structure, achieved through a controlled thermal phase separation process (−80□°C to 4□°C), confirms the effectiveness of the fabrication method in generating an interconnected pore network without phase separation or VC crystallization. Such structural integrity is critical, as it supports nutrient diffusion, cell infiltration, and vascular ingrowth, key factors for successful tissue regeneration.

Unlike conventional HyCol composites that often rely on chemical crosslinkers, potentially compromising biocompatibility, these scaffolds maintain their shape and mechanical stability post-lyophilization, making them suitable for in vivo handling and application. The synergistic combination of hyaluronic acid and collagen provides a biomimetic framework, while the incorporation of vitamin C enhances the scaffold’s therapeutic potential. Collectively, these features underscore the scaffold’s promise as a multifunctional platform for wound healing, offering both long-term stability and bioactive functionality.

### Morphological and Chemical Characterization of Scaffolds

To evaluate the microstructural and chemical properties of the fabricated scaffolds, scanning electron microscopy (SEM) and attenuated total reflectance–Fourier transform infrared spectroscopy (ATR-FTIR) analyses were performed, as shown in **Figure 2**. The SEM images (**Figure 2**, left panel) reveal the porous architecture of all scaffold formulations. The base HyCol scaffold (**A**) exhibits a well-defined, interconnected porous network, which is essential for facilitating cell infiltration, nutrient diffusion, and waste removal. Upon incorporation of vitamin C at increasing concentrations (VC-HyCol-1, −2, and −3; **B–D**), the overall porous morphology is retained, although subtle differences in pore size and wall thickness are observed. Notably, VC-HyCol-3 (**D**), containing the highest vitamin C concentration, displays slightly denser pore walls, which may be attributed to enhanced crosslinking or molecular interactions between vitamin C and the scaffold matrix. These morphological features suggest that the scaffolds maintain structural integrity while allowing tunability based on bioactive loading. The ATR-FTIR spectra (**Figure 2**, right panel) further confirm the successful integration of scaffold components. Characteristic absorption bands for collagen are observed around 1650 cm^−1^ (amide I), 1550 cm^−1^ (amide II), and 1240 cm^−1^ (amide III), indicating the preservation of its triple-helical structure.

**Figure 2.**
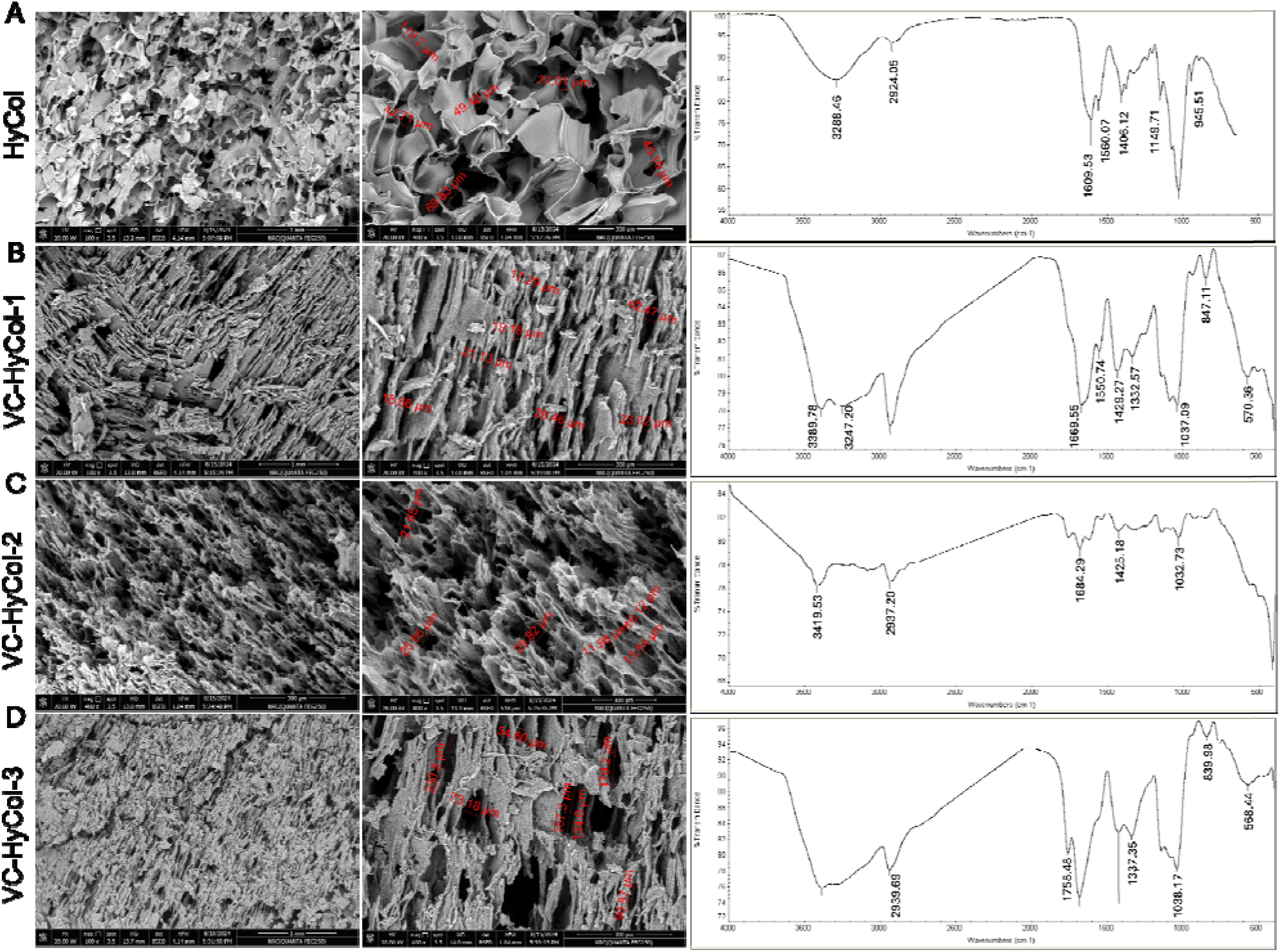
Characterization of all produced scaffolds. Scanning electron microscopy images showing the morphology of the scaffold texture (left). ATR-FTIR data of all scaffolds (right).

Hyaluronic acid contributes to peaks near 1030 cm^−1^, corresponding to C–O–C stretching vibrations. With the addition of vitamin C, new or intensified bands appear in the range of 3200–3500 cm^−1^, associated with O–H stretching, and around 1740 cm^−1^, indicative of C=O stretching from ascorbiccid. These spectral shifts confirm the presence of vitamin C within the scaffold matrix and suggest potential hydrogen bonding interactions that may influence scaffold stability and degradation behavior. Toge her, the SEM and ATR-FTIR analyses validate the successful fabrication of structurally robust and chemically integrated VC-HyCol scaffolds. The preserved porosity and confirmed molecular composition support their suitability for subsequent biological evaluations.

### Physicochemical and Biological Evaluation of VC-HyCol Scaffolds

#### Porosity and Swelling Behavior

As shown in **Figure 3A**, all scaffolds exhibited high porosity, a critical parameter for effective wound healing. The base HyCol scaffold demonstrated substantial porosity, which was slightly reduced with increasing vitamin C content in VC-HyCol-1, −2, and −3. This trend may be attributed to the denser matrix formation due to vitamin C-induced crosslinking or molecular interactions. Nevertheless, all formulations maintained porosity levels conducive to cell infiltration and nutrient exchange. Swelling kinetics, presented in **Figure 3B**, indicate a corresponding increase in swelling capacity with higher VC incorporation. VC-HyCol-3 showed the greatest swelling percentage, which may be attributed to its higher hydrophilic content and more open structure, facilitating greater water uptake. All scaffolds retained sufficient swelling capacity to maintain a moist wound environment, which is essential for optimal healing.

**Figure 3.**
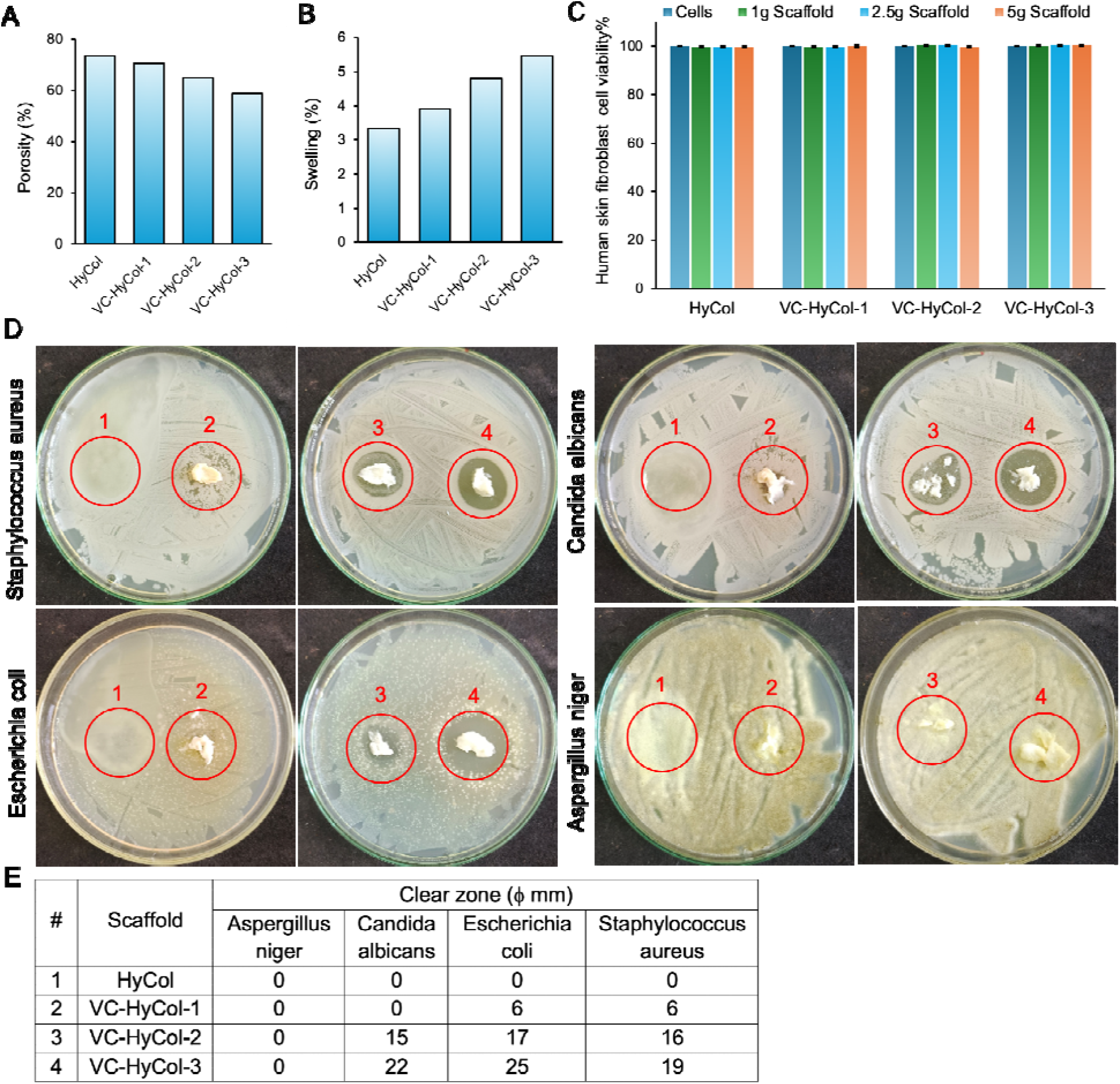
Physicochemical and biological characterization of HyCol scaffolds. (A) Porosity measurements, (B)Swelling kinetics, (C) *In vitro* cytotoxicity against human skin fibroblasts (MTT assay), and (D) Qualitative antibacterial activity (zone of inhibition assay) for: (1) HyCol control, (2) VC-HyCol-1, (3) VC-HyCol-2, and (4) VC-HyCol-3. (E) Quantitative antimicrobial efficacy against clinically relevant pathogens: *Staphylococcus aureus* (Gram-positive), *Escherichia coli* (Gram-negative), *Candida albicans* (yeast), and *Aspergillus niger* (filamentous fungus), presented as inhibition zone diameter (mm).

#### Cytocompatibility

The *in vitro* cytotoxicity assay (MTT) against human skin fibroblasts (**Figure 3C**) confirmed the biocompatibility of all scaffold formulations. Cell viability remained about 100% across all tested scaffold weights (1, 2.5, and 5 g), with no significant cytotoxic effects observed. This suggests that all scaffolds may promote fibroblast proliferation through their role in collagen synthesis and oxidative stress reduction.

#### Antimicrobial Activity

The qualitative zone of inhibition assay (**Figure 3D**) demonstrated that the VC-HyCol scaffolds exhibited clear antimicrobial activity against a range of clinically relevant pathogens, including *Staphylococcus aureus, Escherichia coli, Candida albicans*, and *Aspergillus niger*. In contrast, the HyCol control scaffold showed minimal to no inhibition, particularly against fungal strains. Quantitative analysis (**Figure 3E**) further confirmed these findings. VC-HyCol-2 and VC-HyCol-3 displayed the largest inhibition zones, particularly against *E. coli* (25 mm) and *C. albicans* (22 mm), indicating potent broad-spectrum antimicrobial efficacy. Notably, none of the scaffolds inhibited *Aspergillus niger*, suggesting that additional antifungal agents may be required for applications targeting filamentous fungi.

These results collectively demonstrate that the incorporation of vitamin C enhances not only the structural and swelling properties of the scaffolds but also their biological performance, including cytocompatibility and antimicrobial activity. The VC-HyCol-3 scaffold, in particular, emerged as the most promising formulation, balancing porosity, biocompatibility, and pathogen inhibition.

### *In Vivo* Wound Healing Efficacy in Rat Models

The therapeutic potential of the scaffolds was further evaluated *in vivo* using a full-thickness excisional wound model in rats. **Figure 4** presents both qualitative and quantitative assessments of wound healing progression across different treatment groups over 14 days. **Figure 4** (top panel) displays representative images of wound sites at days 0, 3, 7, 10, and 14 for each group: untreated negative control, HyCol, VC-HyCol-1, VC-HyCol-2, and VC-HyCol-3. While all groups showed gradual wound closure over time,the VC-HyCol-treated groups, particularly VC-HyCol-3, exhibited visibly faster and more com lete healing. By day 14, wounds treated with VC-HyCol-3 were nearly closed, with minimal scarring and well-formed epithelial tissue, in contrast to the control and HyCol groups, which still showed incomplete closure and residual inflammation. Quantitative analysis of wound contraction is shown in **Figure 4** (bottom panel). The percentage of wound closure was significantly higher in all VC-HyCol groups compared to the control and HyCol groups at each time point. Notably, VC-HyCol-3 achieved the highest contraction rates by day 14. These findings confirm that the incorporation of vitamin C into the HyCol scaffold markedly enhances wound healing kinetics. The improved outcomes are likely due to the synergistic effects of structural support from the scaffold matrix, antioxidant activity of vitamin C, and its role in promoting collagen synthesis and angiogenesis. The superior performance of VC-HyCol-3 underscores the importance of optimizing vitamin C concentration for maximal therapeutic benefit.

**Figure 4.**
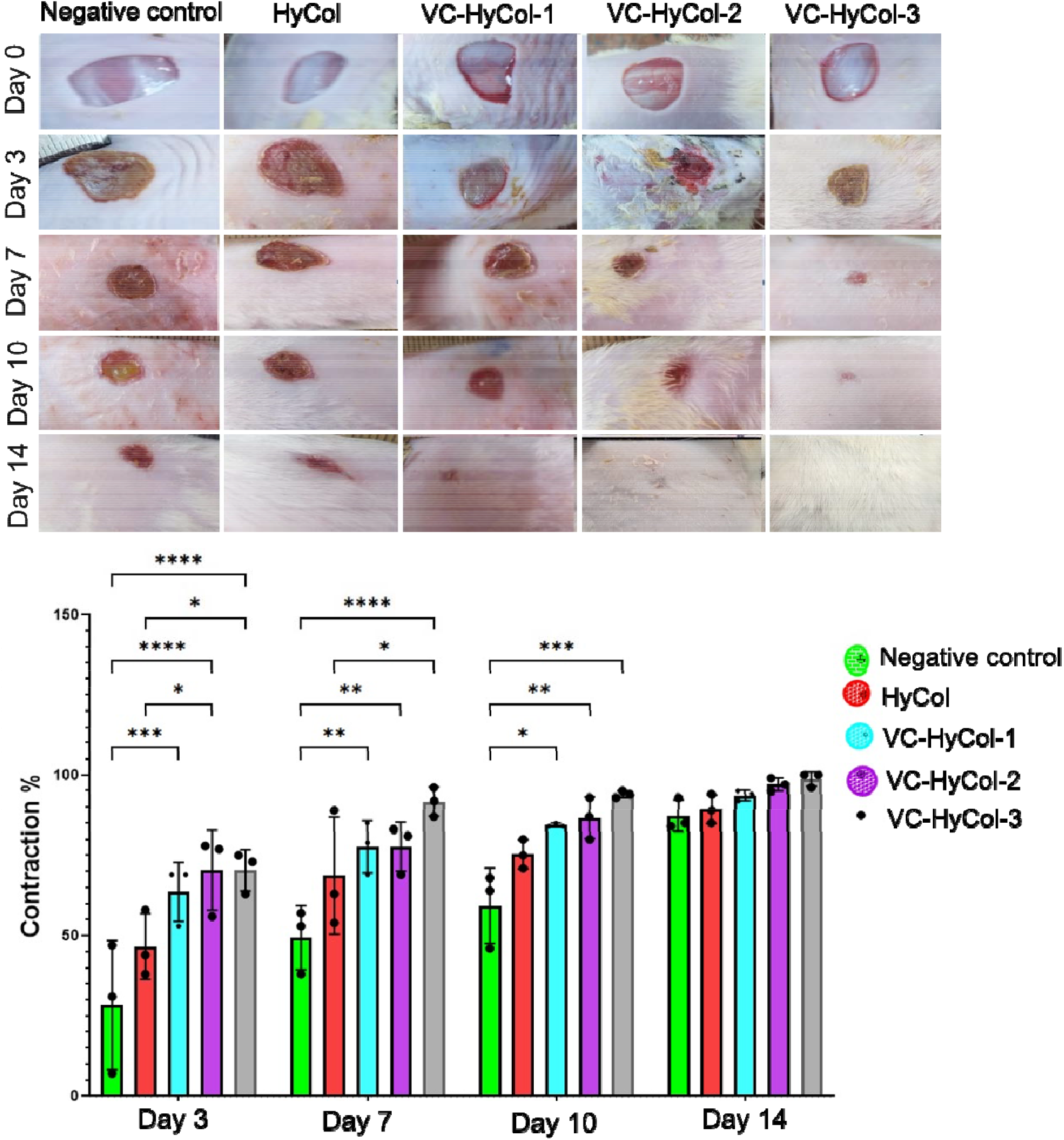
*In vivo* wound healing assessment in rat models treated with different scaffold formulations. **Top:** Representative images of wound sites at Days 0, 3, 7, 10, and 14 for each treatment group: Negative control, HyCol, VC-HyCol-1, VC-HyCol-2, and VC-HyCol-3. **Bottom:** Quantitative analysis of wound contraction (%) over time. Data are presented as mean ± SD. Statistical significance between groups is indicated (*\*p* < 0.05, *\*\*p* < 0.01, *\*\*\*p* < 0.001, *\*\*\*\*p* < 0.0001).

### Biochemical and Molecular Biomarker Analysis

To elucidate the molecular mechanisms underlying the enhanced wound healing observed with VC-HyCol scaffolds, ELISA and western blot analyses were performed to quantify key biomarkers associated with inflammation, oxidative stress, angiogenesis, and tissue regeneration across six scaffolds (**Figure 5A–G**). *Anti-apoptotic and angiogenic markers:* As shown in **Figure 5A–B**, levels of Bcl-2 (an anti-apoptotic protein) and VEGF (vascular endothelial growth factor) were significantly elevated in the VC-HyCol-treated groups, particularly in VC-HyCol-3. This suggests enhanced cell survival and neovascularization, both of which are critical for effective wound repair. The increase in VEGF aligns with the observed histological evidence of neovascularization and supports the scaffold’s pro-angiogenic potential. *Oxidative Stress and Antioxidant Defense:* Glutathione (GSH), a key antioxidant, was significantly upregulated in VC-HyCol groups (**Figure 5C**), while malondialdehyde (MDA), a marker of lipid peroxidation and oxidative damage, was markedly reduced (**Figure 5G**). These findings confirm the antioxidant role of vitamin C within the scaffold, effectively mitigating oxidative stress in the wound microenvironment. *Inflammatory Cytokines:* Pro-inflammatory cytokines TNF-α and IL-1β were significantly downregulated in VC-HyCol-treated groups compared to controls (**Figure 5D and 5F**), indicating a strong anti-inflammatory effect. This is further supported by the reduced expression of NF-κB p65, a central transcription factor in inflammatory signaling, as shown in the western blot (**Figure 5H**) and its corresponding densitometric analysis (**Figure 5I**). The suppression of NF-κB activity suggests that the scaffold modulates inflammatory pathways at the transcriptional level. *Angiogenesis and Stem Cell Marker:* CD34, a marker associated with endothelial progenitor cells and neovascularization, was significantly elevated in VC-HyCol-2 and VC-HyCol-3 (**Figure 5E**), reinforcing the scaffold’s role in promoting vascular regeneration. Collectively, these molecular findings provide mechanistic insight into the scaffold’s therapeutic efficacy. The VC-HyCol-3 scaffold, in particular, demonstrated the most favorable biomarker profile, enhancing antioxidant defenses, promoting angiogenesis, and suppressing inflammation, thereby creating an optimal environment for tissue regeneration.

**Figure 5.**
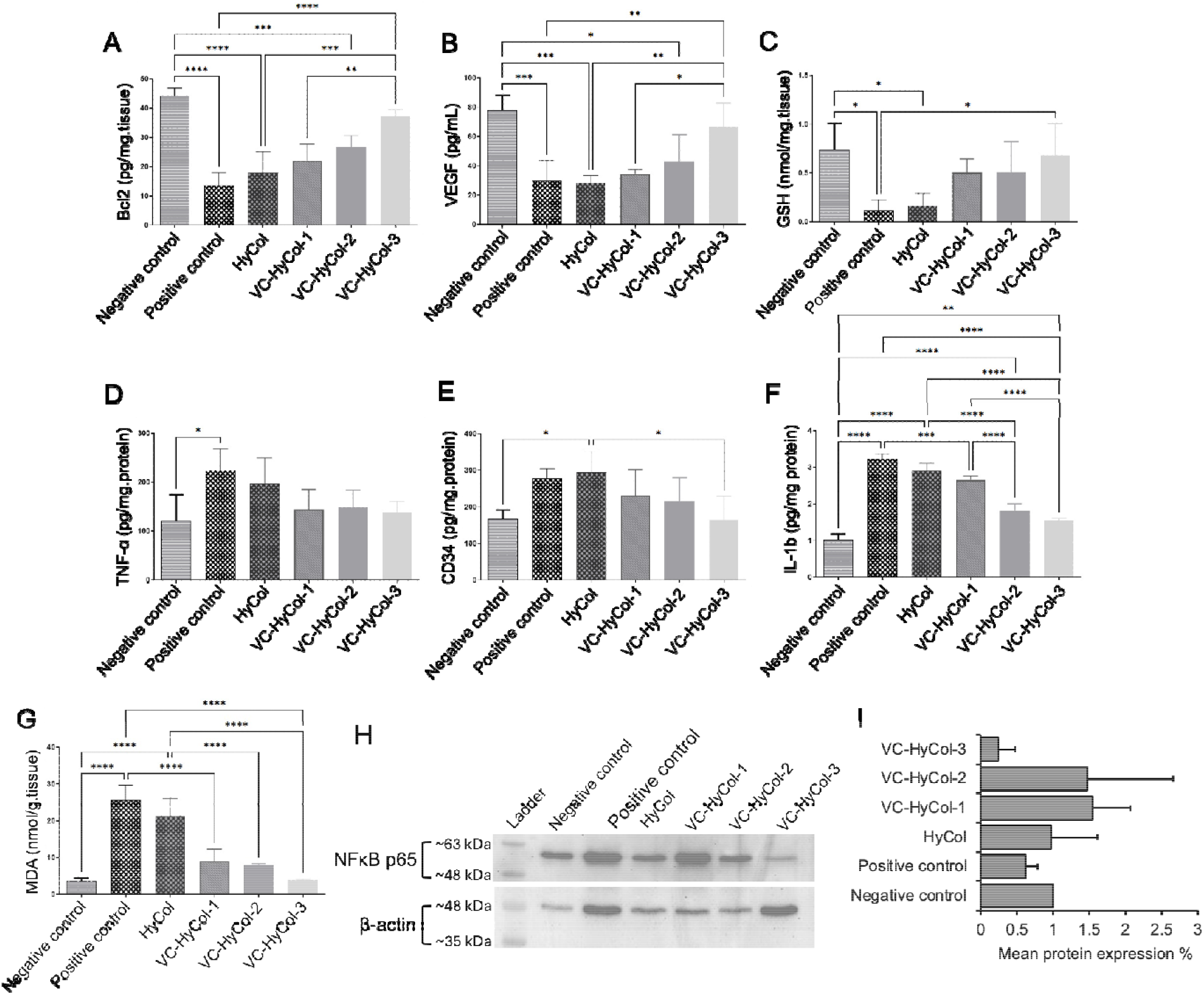
Molecular analysis of inflammatory, oxidative, and regenerative biomarkers in rat wound tissues following treatment with different scaffold formulations. **(A–G):** ELISA quantification of Bcl-2, VEGF, GSH, TNF-α, CD34, IL-1β, and MDA levels across six groups: Negative control, positive control, HyCol, VC-HyCol-1, VC-HyCol-2, and VC-HyCol-3. **(H):** Western blot analysis of NF-κB p65 and β-actin expression. **(I):** Densitometric quantification of NF-κB p65 normalized to β-actin. Data are presented as mean ± SD. Statistical significance is indicated by asterisks (*\*p* < 0.05, *\*\*p* < 0.01, *\*\*\*p* < 0.001, *\*\*\*\*p* < 0.0001).

### Histopathological Evaluation of Wound Healing

Histological analysis was performed to assess tissue regeneration and inflammatory response following treatment with various scaffolds. Representative hematoxylin and eosin (H&E) stained sections are shown in **Figure 6**, highlighting key features such as fibrous connective tissue (II]), mononuclear inflammatory cell infiltration (red arrows), epidermal integrity (black arrows), and edema (blue arrows). In the negative control group (**Figure 6A**), tissue architecture remained largely intact, with a organized epidermis and dermis, minimal inflammatory infiltration, and no observable edema. well-This served as a baseline for healthy tissue morphology. The positive control group (**Figure 6B**), in contrast, exhibited pronounced pathological changes. There was significant disruption of the epidermal layer, extensive infiltration of mononuclear inflammatory cells, and marked edema, as evidenced by widened interstitial spaces and separation of dermal fibers. The fibrous connective tissue appeared disorganized and degraded, indicating impaired healing. Treatment with the HyCol scaffold (**Figure 6C**) led to partial restoration of tissue structure. The epidermis showed improved continuity, and inflammatory infiltration was reduced compared to the positive control. However, moderate edema and irregular fibrous tissue organization persisted, suggesting limited regenerative capacity. The VC-HyCol-1 group (**Figure 6D**) demonstrated further histological improvement. The epidermis was largely intact, with minimal inflammatory cell presence. Edema was less pronounced, and fibrous connective tissue appeared more structured, indicating enhanced tissue remodeling. In the VC-HyCol-2 group (**Figure 6E**), histolo ical features closely resembled those of healthy tissue. The epidermis was continuous and well-formed, inflammatory infiltration was minimal, and edema was substantially reduced. Fibrous connective tissue was densely packed and well-organized, reflecting effective extracellular matrix remodeling and wound resolution. The VC-HyCol-3 group (**Figure 6F**) exhibited the most favorable histological profile. Epidermal integrity was fully restored, inflammatory cells were nearly absent, and edema was completely resolved. The fibrous connective tissue was indistinguishable from that of the negative control, indicating complete tissue regeneration. Overall, the histopathological evidence supports the superior healing efficacy of VC-HyCol-3, which facilitated near-complete tissue restoration with minimal inflammatory response, pro-regenerative and structural remodeling properties highlighting its potential as an advanced wound dressing.

**Figure 6.**
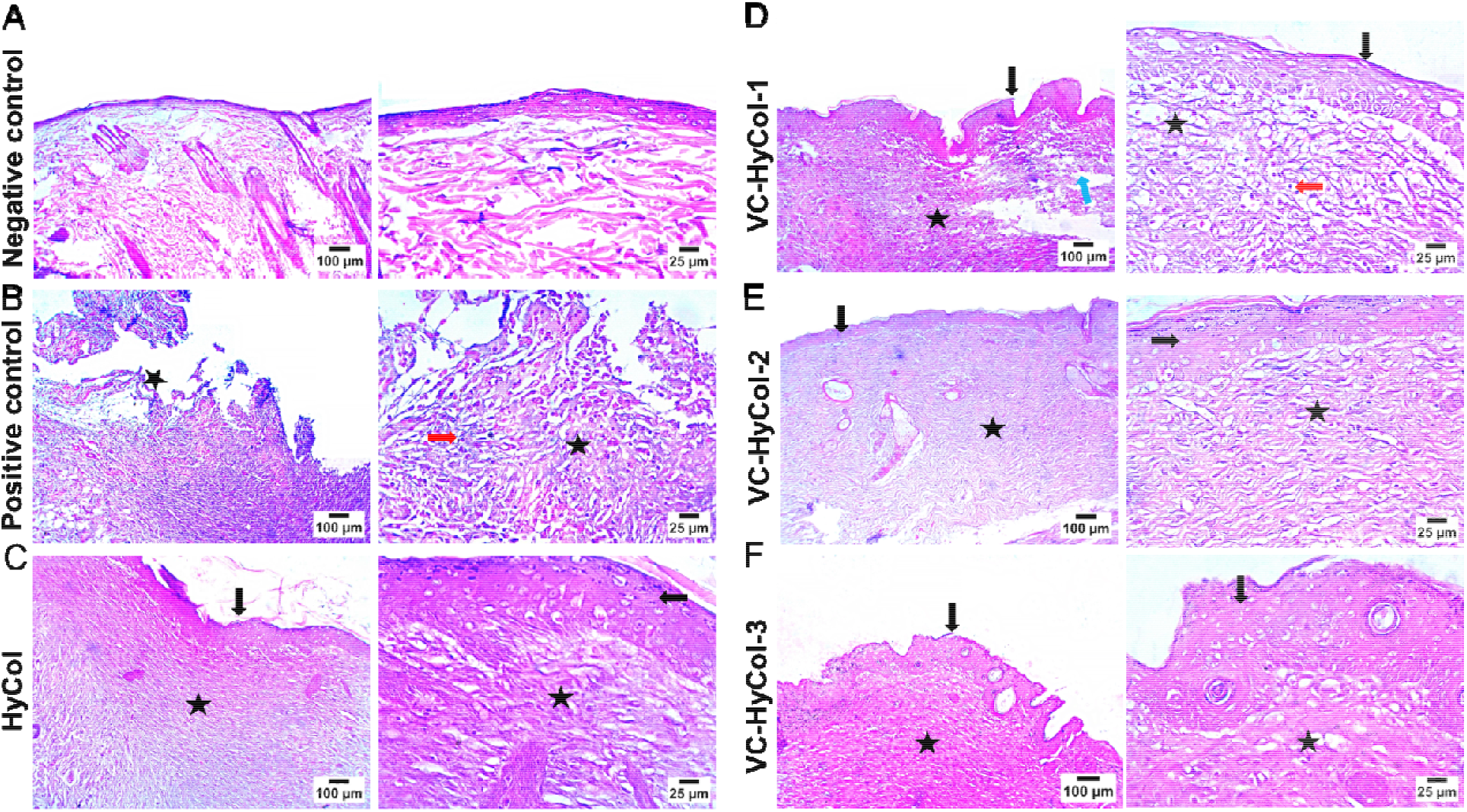
Histopathological analysis of wound tissue sections stained with hematoxylin and eosin (H&E) from different treatment groups on day 14 post-injury. (A) Negative control showing normal skin architecture. (B) Positive control with disrupted epidermis (black arrow), fibrous connective tissue (star), and mononuclear inflammatory cell infiltration (red arrow). (C) HyCol group with partially restored epidermis and connective tissue. (D) VC-HyCol-1 showing improved epidermis and connective tissue, with residual inflammation (red arrow) and edema (blue arrow). (E) VC-HyCol-2 and (F) VC-HyCol-3 exhibiting intact epidermis, organized fibrous tissue, and minimal inflammation or edema. These results confirm enhanced tissue regeneration and reduced inflammation in vitamin C-loaded scaffolds, particularly VC-HyCol-3.

### Immunohistochemical Analysis of Tissue Regeneration

To further evaluate the biological activity of the scaffolds at the cellular level, immunohistochemistry (IHC) was performed to assess the expression of specific markers associated with tissue regeneration and remodeling. Representative stained tissue sections from each treatment group are shown in **Figure 7A–F**, with quantitative analysis summarized in **Figure 7G. Figure 7A–F** shows immunostained sections from the negative control, positive control, HyCol, VC-HyCol-1, VC-HyCol-2, and VC-HyCol-3 groups, respectively. The red arrows highlight regions of positive staining, indicating the presence of target proteins associated with tissue regeneration and remodeling. The negative control group (**Figure 7A**) exhibited minimal staining, while the positive control (**Figure 7B**) showed moderate expression, likely due to natural healing processes. The HyCol group (**Figure 7C**) demonstrated increased staining intensity, suggesting some regenerative activity. However, the VC-HyCol-treated groups (**Figure 7D–F**) showed markedly enhanced staining, with VC-HyCol-3 (**Figure 7F**) displaying the most extensive and intense signal, indicating robust expression of regenerative markers. This suggests that the scaffold not only supports structural repair but also actively promotes cellular processes involved in tissue regeneration. Quantitative analysis of the immunoreactive area (**Figure 7G**) confirmed these observations. VC-HyCol-3 exhibited the highest percentage of positive staining, significantly surpassing all other groups (*p* < 0.001), followed by VC-HyCol-2 and VC-HyCol-1. These results align with previous histological and molecular findings, reinforcing the conclusion that vitamin C-loaded scaffolds, particularly at higher concentrations, enhance the biological activity of the wound bed, accelerating and improving the quality of healing.

**Figure 7.**
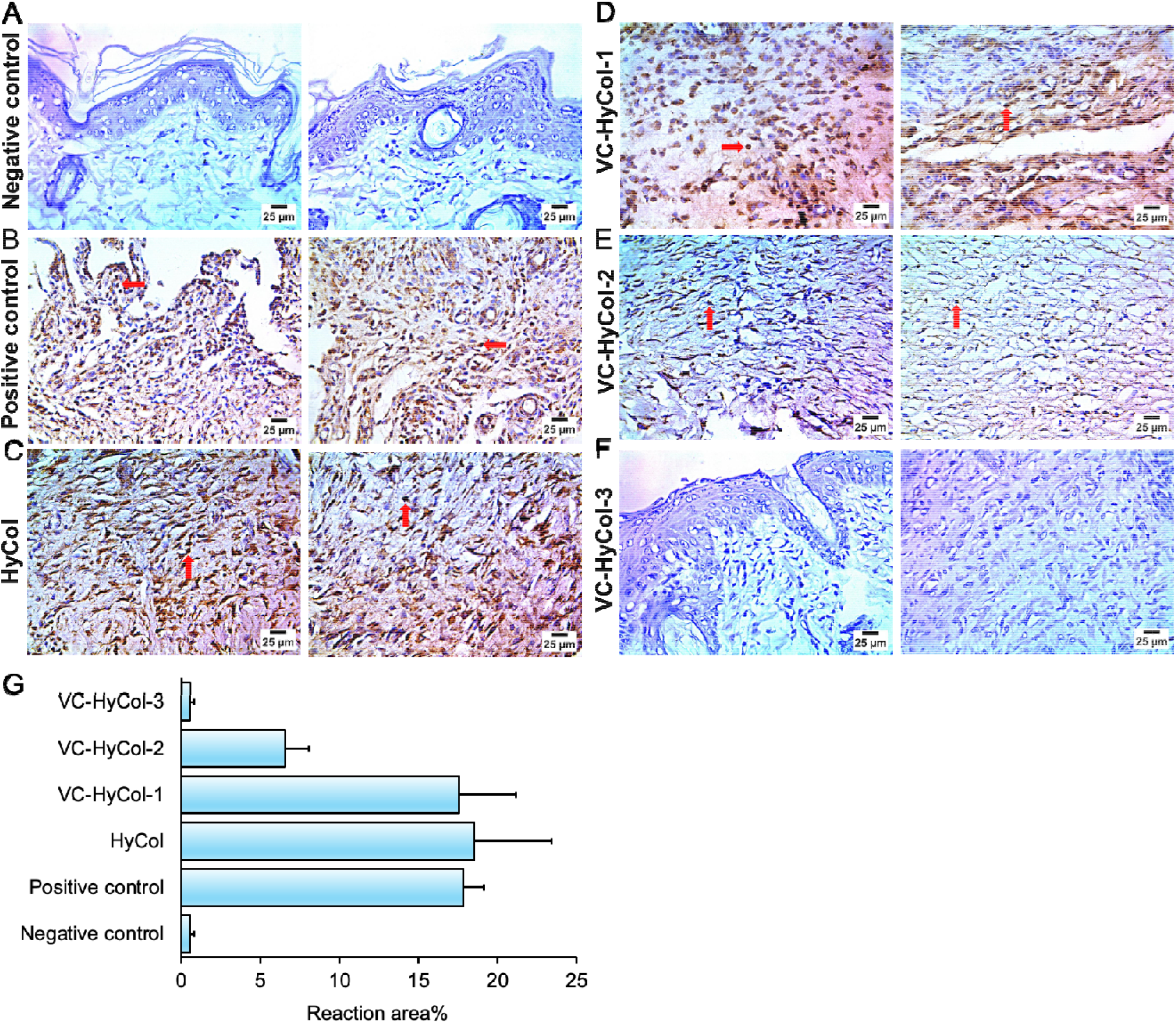
Immunohistochemical analysis of wound tissue sections from different treatment groups. (A–F) Representative IHC-stained images from negative control, positive control, HyCol, VC-HyCol-1, VC-HyCol-2, and VC-HyCol-3 groups. Red arrows indicate regions of positive staining. Scale bars = 25 µm. (G) Quantitative analysis of the immunoreactive area (%), showing significantly increased marker expression in VC-HyCol-treated groups, especially VC-HyCol-3. Data are presented as mean ± SD. Statistical significance: *\*p* < 0.05, *\*\*p* < 0.01, *\*\*\*p* < 0.001.

## Conclusion

This study successfully developed and evaluated a novel vitamin C-loaded hyaluronic acid–collagen (VC-HyCol) scaffold designed to enhance wound healing through synergistic structural, biochemical, and antioxidant mechanisms. The one-pot fabrication method yielded scaffolds with favorable porosity, mechanical integrity, and sustained vitamin C incorporation, as confirmed by SEM and ATR-FTIR analyses. Physicochemical assessments demonstrated that the scaffolds maintained high porosity and swelling capacity, essential for supporting a moist wound environment and cellular infiltration. Importantly, the VC-HyCol scaffolds exhibited excellent cytocompatibility and potent antimicrobial activity, particularly against *E. coli* and *C. albicans*, with VC-HyCol-3 showing the most pronounced effects. *In vivo* studies in a rat full-thickness wound model revealed that VC-HyCol-3 significantly accelerated wound closure, reduced inflammation, and promoted organized tissue regeneration. ELISA and western blot analyses further confirmed that the scaffold modulated key molecular pathways by enhancing antioxidant defenses (↑GSH, ↓MDA), promoting angiogenesis (↑VEGF, ↑CD34), and suppressing inflammatory mediators (↓TNF-α, ↓IL-1β, ↓NF-κB p65). Histological and immunohistochemical evaluations corroborated these findings, showing restored epidermal architecture, reduced edema and inflammatory infiltration, and elevated expression of regenerative markers in VC-HyCol-3-treated wounds. Collectively, these results highlight the therapeutic potential of VC-HyCol scaffolds, particularly the VC-HyCol-3 formulation, as a multifunctional wound dressing capable of accelerating healing through structural support, oxidative stress mitigation, and immunomodulation. This platform offers a promising, clinically translatable strategy for managing chronic and hard-to-heal wounds. Future studies may explore its application in diabetic ulcers, burn injuries, and other complex wound environments, as well as its scalability and performance in larger animal models or clinical trials.

## References

[1] C. Wang et al., “Wound management materials and technologies from bench to bedside and beyond,” Nature Reviews Materials, vol. 9, no. 8, pp. 550–566, 2024.

[2] A. Sharma, R. Shankar, A. K. Yadav, A. Pratap, M. A. Ansari, and V. Srivastava, “Burden of chronic nonhealing wounds: an overview of the worldwide humanistic and economic burden to the healthcare system,” The International Journal of Lower Extremity Wounds, p. 15347346241246339, 2024.

[3] S. G. Pereira, J. Moura, E. Carvalho, and N. Empadinhas, “Microbiota of chronic diabetic wounds: ecology, impact, and potential for innovative treatment strategies,” Frontiers in microbiology, vol. 8, p. 296236, 2017.

[4] M. S. Brown, B. Ashley, and A. Koh, “Wearable technology for chronic wound monitoring: current dressings, advancements, and future prospects,” Frontiers in bioengineering and biotechnology, vol. 6, p. 47, 2018.

[5] Y. Cai, K. Chen, C. Liu, and X. Qu, “Harnessing strategies for enhancing diabetic wound healing from the perspective of spatial inflammation patterns,” Bioactive materials, vol. 28, pp. 243–254, 2023.

[6] P. J. Muire, M. A. Thompson, R. J. Christy, and S. Natesan, “Advances in immunomodulation and immune engineering approaches to improve healing of extremity wounds,” International Journal of Molecular Sciences, vol. 23, no. 8, p. 4074, 2022.

[7] M. Hunt, M. Torres, E. Bachar-Wikström, and J. D. Wikström, “Multifaceted roles of mitochondria in wound healing and chronic wound pathogenesis,” Frontiers in Cell and Developmental Biology, vol. 11, p. 1252318, 2023.

[8] H. Yu et al., “Landscape of the epigenetic regulation in wound healing,” Frontiers in Physiology, vol. 13, p. 949498, 2022.

[9] L. Uccioli, V. Izzo, M. Meloni, E. Vainieri, V. Ruotolo, and L. Giurato, “Non-healing foot ulcers in diabetic patients: general and local interfering conditions and management options with advanced wound dressings,” Journal of wound care, vol. 24, no. Sup4b, pp. 35–42, 2015.

[10] J. Dawi et al., “Diabetic Foot Ulcers: Pathophysiology, Immune Dysregulation, and Emerging Therapeutic Strategies,” Biomedicines, vol. 13, no. 5, p. 1076, 2025.

[11] S. Dhivya, V. V. Padma, and E. Santhini, “Wound dressings–a review,” BioMedicine, vol. 5, no. 4, p. 22, 2015.

[12] J. Boateng and O. Catanzano, “Advanced therapeutic dressings for effective wound healing—a review,” Journal of pharmaceutical sciences, vol. 104, no. 11, pp. 3653–3680, 2015.

[13] Y. Zhao, Y. Zhao, B. Xu, H. Liu, and Q. Chang, “Microenvironmental dynamics of diabetic wounds and insights for hydrogel-based therapeutics,” Journal of Tissue Engineering, vol. 15, p. 20417314241253290, 2024.

[14] J.-D. Xue, J. Gao, A.-F. Tang, and C. Feng, “Shaping the immune landscape: Multidimensional environmental stimuli refine macrophage polarization and foster revolutionary approaches in tissue regeneration,” Heliyon, vol. 10, no. 17, 2024.

[15] A. Salerno and P. A. Netti, “Review on bioinspired design of ECM-mimicking scaffolds by computer-aided assembly of cell-free and cell laden micro-modules,” Journal of Functional Biomaterials, vol. 14, no. 2, p. 101, 2023.

[16] F. Urciuolo, G. Imparato, and P. Netti, “In vitro strategies for mimicking dynamic cell–ECM reciprocity in 3D culture models,” Frontiers in Bioengineering and Biotechnology, vol. 11, p. 1197075, 2023.

[17] Y. Chen, X. Dong, M. Shafiq, G. Myles, N. Radacsi, and X. Mo, “Recent advancements on three-dimensional electrospun nanofiber scaffolds for tissue engineering,” Advanced Fiber Materials, vol. 4, no. 5, pp. 959–986, 2022.

[18] S. Oh, S. B. Seo, G. Kim, S. Batsukh, K. H. Son, and K. Byun, “Poly-D, L-lactic acid stimulates angiogenesis and collagen synthesis in aged animal skin,” International Journal of Molecular Sciences, vol. 24, no. 9, p. 7986, 2023.

[19] H. Yang et al., “Multifunctional hydrogel targeting senescence to accelerate diabetic wound healing through promoting angiogenesis,” Journal of Nanobiotechnology, vol. 23, no. 1, p. 177, 2025.

[20] E. S. Armengol et al., “Unveiling the potential of biomaterials and their synergistic fusion in tissue engineering,” European Journal of Pharmaceutical Sciences, p. 106761, 2024.

[21] S. M. Riha, M. Maarof, and M. B. Fauzi, “Synergistic effect of biomaterial and stem cell for skin tissue engineering in cutaneous wound healing: A concise review,” Polymers, vol. 13, no. 10, p. 1546, 2021.

[22] J. P. Rodrigues et al., “Development of collagenous scaffolds for wound healing: characterization and in vivo analysis,” Journal of Materials Science: Materials in Medicine, vol. 35, no. 1, p. 12, 2024.

[23] Y. Dong and Z. Wang, “ROS-scavenging materials for skin wound healing: advancements and applications,” Frontiers in Bioengineering and Biotechnology, vol. 11, p. 1304835, 2023.

[24] Q. Yu et al., “Effects of reactive oxygen species and antioxidant strategies on wound healing in diabetes,” Interdisciplinary Medicine, vol. 3, no. 2, p. e20240062, 2025.

[25] H. Sarpooshi, M. Haddadi, M. Siavoshi, and R. Borghabani, “Wound healing with Vitamin C,” Transl Biomed, vol. 8, no. 4, pp. 139–42, 2017.

[26] N. Bechara, V. Flood, and J. Gunton, “A systematic review on the role of vitamin C in tissue healing. Antioxidants. 2022; 11 (8): 1605,” ed.

[27] A. S. Montaser, K. Jlassi, M. A. Ramadan, A. A. Sleem, and M. F. Attia, “Alginate, gelatin, and carboxymethyl cellulose coated nonwoven fabrics containing antimicrobial AgNPs for skin wound healing in rats,” International Journal of Biological Macromolecules, vol. 173, pp. 203–210, 2021.

[28] M. F. Attia, R. Akasov, F. Alexis, and D. C. Whitehead, “Polymer-scaffolded synthesis of periodic mesoporous organosilica nanomaterials for delivery systems in cancer cells,” ACS Biomaterials Science & Engineering, vol. 6, no. 12, pp. 6671–6679, 2020.

[29] N. Percie du Sert et al., “The ARRIVE guidelines 2.0: Updated guidelines for reporting animal research,” Journal of Cerebral Blood Flow & Metabolism, vol. 40, no. 9, pp. 1769–1777, 2020.

[30] Z. Xiao et al., “Pharmacological effects of salvianolic acid B against oxidative damage,” Frontiers in Pharmacology, vol. 11, p. 572373, 2020.

[31] A. A. Sedik, M. Salama, K. Fathy, and A. Salama, “Cold plasma approach fortifies the topical application of thymoquinone intended for wound healing via up-regulating the levels of TGF-ß, VEGF, and α-SMA in rats,” International Immunopharmacology, vol. 122, p. 110634, 2023.

[32] G. A. Ujah et al., “Protective effect of tert-butylhydroquinone against cisplatin-induced hepatorenal injury via modulating oxidative stress, inflammation, and apoptosis,” Archives of Physiology and Biochemistry, vol. 130, no. 6, pp. 951–961, 2024.

[33] Y. Zhao et al., “Salvianolic acid B inhibits atherosclerosis and TNF-α-induced inflammation by regulating NF-κB/NLRP3 signaling pathway,” Phytomedicine, vol. 119, p. 155002, 2023.

[34] F. Lv et al., “A conducive bioceramic/polymer composite biomaterial for diabetic wound healing,” Acta biomaterialia, vol. 60, pp. 128–143, 2017.

